# Time course of lipopolysaccharide- and capsaicin-induced cytokine release in an ex vivo mouse model of vaginal inflammation

**DOI:** 10.1101/2023.02.11.528146

**Authors:** K Jane Chalmers, Mark R Hutchinson, Kelsi N Dodds, Yuen H Kwok, Susan F Evans, G Lorimer Moseley

## Abstract

**Problem:** The neuroimmune interface has been characterised in few areas of the body. The objective of this study was to investigate the neuroimmune interface in the mouse vagina through a novel *ex vivo* model, to determine whether LPS could directly activate and produce TRPV1-mediated neuronal activation.

**Method of Study:** Concentrations of IL-1β and IL-6 release into the supernatant at different times post *ex vivo* stimulation with LPS, capsaicin, or a combination of the two were assessed using enzyme-linked immunosorbent assay.

**Results:** There were no differences in the supernatant concentration of IL-6 with different stimulation type nor stimulation time. Supernatant concentrations of IL-1β were greater at the 20 hour time point than the 4 hour time point, and were greater for stimulations involving LPS.

**Conclusion:** There is a clear pattern of pro-inflammatory neuroimmune responses following *ex vivo* stimulation of mouse vaginal tissues with capsaicin and LPS, evident as an increased IL-1β output. This output is greatest at 20 hours post-stimulation, indicating this neuroimmune response is time-dependent.

## Introduction

The neuroimmune interface has received growing attention for its role in pathological pain conditions^1^. This interface can be thought of as the signalling between the immune and immune-like cells that reside in the central nervous system, and neuronal projections. Neuronal excitability can be enhanced by pro-inflammatory immune mediators released from immunocompetent cells, and conversely, activation of neurons by certain stimuli can initiate pro-inflammatory immune signalling^2,3^. A similar neuroimmune interface can also be found peripherally, albeit with differing immune cell and neuronal types. Peripherally, the neuroimmune interface is integral to pathophysiological changes at sites of tissue injury and to inflammation-induced peripheral sensitization. For example, several immune cells including macrophages^4^, mast cells^5^, and neutrophils^6^ play a role in the periphery in the development of hyperalgesia around a peripheral nerve injury. In parallel, microglia become reactive at the level of spinal innervation^7^, undergo reactive gliosis, and release a host of pro-inflammatory mediators^8^. These mediators allow the migration of other immune cells to the area^9^ as well as sensitising nearby neurons^10^. The release of these mediators also provides a feedforward loop in which further immune cells are activated and migrate to the area^3^, eventually leading to a spreading nociceptive hypersensitivity that is characteristic of, and associated with, central sensitization.

Toll-Like Receptor 4 (TLR4) has been identified as a key player in neuroimmune interactions. Many immune cells such as microglia, astrocytes, mast cells, and endothelial cells express TLR4 and the obligatory co-receptor and accessory signalling proteins. However, the expression of TLR4 has also been demonstrated on neurons following injury^11,12^. Thus, TLR4 has an ability to influence both the neuronal and immune systems. As such, TLR4 has been the subject of much research investigating the role of neuroimmune interactions in many conditions such as sepsis^13^ and chronic pain^14^.

Pre-clinical models have aimed to simulate the neuroimmune interaction in chronic pain through pharmacological methods. One example is the co-activation of transient receptor potential cation channel subfamily V member 1 (TRPV1) and TLR4, which potentiates TRPV1 activity in sensory neurons. This activation of the pain-associated neuroimmune signalling has been demonstrated *in vivo* in both animal^15^ and human^16^ models using two stimulants: lipopolysaccharide (LPS) and capsaicin. LPS is a large molecule found in the outer membrane of Gram-negative bacteria^17^. While LPS can be derived from a host of bacteria, the mechanism of activating the host immune system is consistent: through activation of TLR4 on the surface of immunocompetent cells^17^. Capsaicin is a compound that activates sensory neurons and causes the release of vasoactive and pro-inflammatory peptides such as substance P^18^ and CGRP^19^. Capsaicin activates sensory neurons via TRPV1 channels^20^.

Owing to the translational capacity of the endotoxin plus capsaicin stimuli to model exaggerated pain states, several *in vivo* preclinical and clinical studies have been performed. However, there are limitations to this approach. Intravenous endotoxin (LPS) can be accompanied by significant and potentially harmful side effects such as fever^21^, diarrhea^21^, vomiting^21^, hypotension^21^, and septic shock^22^, and these may lead to an unblinding of the subject to the treatment. Furthermore, the *in-vivo* method makes it difficult to exclude system-wide influences on the experimental outcome, such as learning and contextual variables.

An alternative approach to indirectly assessing the neuroimmune interface is to engage these classic neuronal and immunological signals outside of the body. This method enables an investigation into the neuroimmune interface in specific and defined areas, without the influence of external system-wide factors. A recent study using a rat model has demonstrated the ability of TLR4 to potentiate TRPV1 activity in trigeminal ganglia (TG) sensory neurons using LPS and capsaicin stimulations *in vitro*^*15*^. In this study, TG neurons were isolated and used as a primary culture for 5 days of stimulation. This was the first study of its kind to demonstrate a region-specific neuroimmune interaction using *in vitro* methods.

Characterising the neuroimmune interface in different body regions is essential to understand the normal and abnormal function of these areas. An exploration of the neuroimmune interface of the reproductive tract may reveal potential mechanistic drivers for certain conditions in this area where a dysfunctional neuroimmune interface is indicated. For example, vulvodynia, a condition that is characterised by vulvar hyperalgesia, and which exerts substantial psychological, behavioural, and social impact^23^ on around 12% of women over their lifetime^24^. Vulvodynia is associated with previous repeated infections of *Candida albicans*^24^ *and a higher density of TRPV1 positive sensory nerves within vestibular tissues*^*25,26*^. *Animal models have demonstrated that repeated vulvovaginal infections with Candida albicans* can lead to vulvar hyperalgesia^27^. These findings, taken with the fact that *Candida albicans* is known to elicit an immune response through activation of TLR4^28,29^, suggest that vulvodynia may be a condition characterised by a dysfunctional neuroimmune interface.

We were interested in exploring the vagina-specific neuroimmune interface using a novel *ex vivo* mouse model. The mouse vagina, similar to the human vagina, is highly innervated^30^, comprised of stratified squamous epithelial cells^31^, and contains an abundance of immune cells^32-35^. In the present study, we aimed to investigate the characteristics of the neuroimmune interface in the mouse vagina. We hypothesised that LPS would directly activate TLR4 and produce TRPV1 (capsaicin)-mediated neuronal activation. Specifically, we hypothesised that both stimulation with LPS and capsaicin alone would generate a heightened pro-inflammatory response compared to a vehicle, observed as an increase in IL-1β and IL-6 levels, but that stimulation with a combination of LPS and capsaicin would generate the greatest pro-inflammatory response, indicative of TLR4 producing TRPV1-mediated neuronal activation.

## Materials and Methods

### Animals

Twelve virgin female BALB/c mice aged 8-12 weeks were obtained from the University of Adelaide Laboratory Animal Services. Cytological evaluation of vaginal smears from the mice was used to determine their stage of estrous, as previously described^36^. Mice in proestrous were selected, corresponding to the estrous stage of high estrogen^37^. All experimental procedures were performed in accordance with the University of Adelaide Animal Ethics Guidelines.

### Experimental protocol

Mice were anesthetized by isoflurane inhalation and culled by cervical dislocation. The vaginal canal was removed and placed in phosphate-buffered saline (PBS; 0.01M; pH 7.4) in a petri dish. Any remaining connective tissue, fat, urethra, rectum, and cervix were removed. The vagina was opened along the anterior edge and pinned plat to the Sylgard-coated base (Dow Corning; MI, USA) of the petri dish using entomology pins. Any remaining mucous was removed from the interior of the vagina. The vagina was divided into four pieces, blot dried using a Kim-wipe, and weighed.

The four tissue pieces were stimulated with 100 μL with either: vehicle (RPMI supplemented with FBS and penicillin), 25ng/mL LPS, 125nM capsaicin, or 25ng/mL LPS + 125nM capsaicin. The concentrations of LPS^38,39^ and capsaicin^15,40^ were based on previous literature demonstrating the ability of these concentrations to induce a pro-inflammatory response. The tissues were incubated at 37°C, 5%CO_2_ for one of four pre-determined times: either 4, 6, 20, or 24 hours. The tissues from three mice were stimulated for 4 hours, from another three mice for 6 hours, from another three mice for 20 hours, and the final three mice for 24 hours. The time period of 4-24 hours was chosen for two reasons. First, previous research has demonstrated that LPS has the ability to potentiate TRPV1 responses within 5 minutes, and can still be observed 5 days later^15^, so 4 hours was determined a short enough period to observe a response. Second, cells needed to remain viable to produce a response, so the stimulation time was capped at 24 hours in order to maximise cell viability while maintaining an observable response. Following the incubation period, the supernatant and tissue were separated and frozen individually at −80°C until the ELISA analysis was completed.

### Enzyme-linked immunosorbent assays (ELISAs)

The main outcome of the study was the supernatant concentration of IL-1β and IL-6. These two cytokines were analysed as they are pro-inflammatory and have previously been demonstrated to increase after stimulation with LPS and capsaicin^16,39,41^. Concentrations of IL-1β and IL-6 were assayed according to protocol using commercially available mouse IL-1β and IL-6 ELISA (BD Bioscience, Australia). UV absorbance was quantified on a BMG PolarStar microplate reader (BMG Labtechnologies, Offenburg, Germany) at 450nm with absorbance at 570nm subtracted. The ELISAs were performed in duplicate and averaged.

### Data Handling

Data were weighted according to the mass of the original tissue piece to account for variations in the size of these excised tissues. Where the detectable amounts of IL-1β and IL-6 in samples were below the limit of quantification (LOQ), the LOQ was used as the value for these data. The LOQ data was then used in the mass-weighting process. Data were inspected visually to determine outliers on the replicated ELISAs. Where outliers were identified, these were removed.

### Statistical Analysis

All experiments were performed in duplicate. Normality of the data were assessed by the Kolmogorov-Smirnov test. In cases where data were not normally distributed, a log transformation was performed and statistical analysis was then conducted using these data. For each cytokine (IL-1β and IL-6), two separate analyses were undertaken. First, three-way ANOVAs were used to assess the effects of stimulation time, stimulation type, and the interaction of the two. Second, the area under the curve (AUC) for each stimulation was calculated, and two-way ANOVAs on the AUC data were used to assess the effects of stimulation type. Statistical significance was set at p<0.05. The results were analysed with Prism software version 7 (GraphPad Software, San Diego, CA, USA).

## Results

### LPS + capsaicin has no impact on stimulated IL-6 release from vaginal tissue sections

To explore the impact of LPS, capsaicin, and the combination effect of LPS+capsaicin on stimulated vaginal peripheral immune responsivity, the IL-6 release into the supernatant of cultures was measured. A three-way ANOVA of IL-6 release into the supernatant revealed no effect of stimulation time (*F*_3,32_=2.016, p>0.05), LPS (*F*_1,32_=0.06724, p>0.05), capsaicin (*F*_1,32_=1.884, p>0.05), nor an interaction between the three (*F*_3,32_=0.6343, p>0.05)(Figure 1a). In order to integrate the effect of time, an area under the curve (AUC) calculation was performed. The two-way ANOVA of the IL-6 (AUC) data also revealed no effect of LPS (*F*_1,8_=1.74, p>0.05), capsaicin, (*F*_1,8_=0.5165, p>0.05), nor an interaction between the two (*F*_1,8_=0.6755, p>0.05)(Figure 2a).

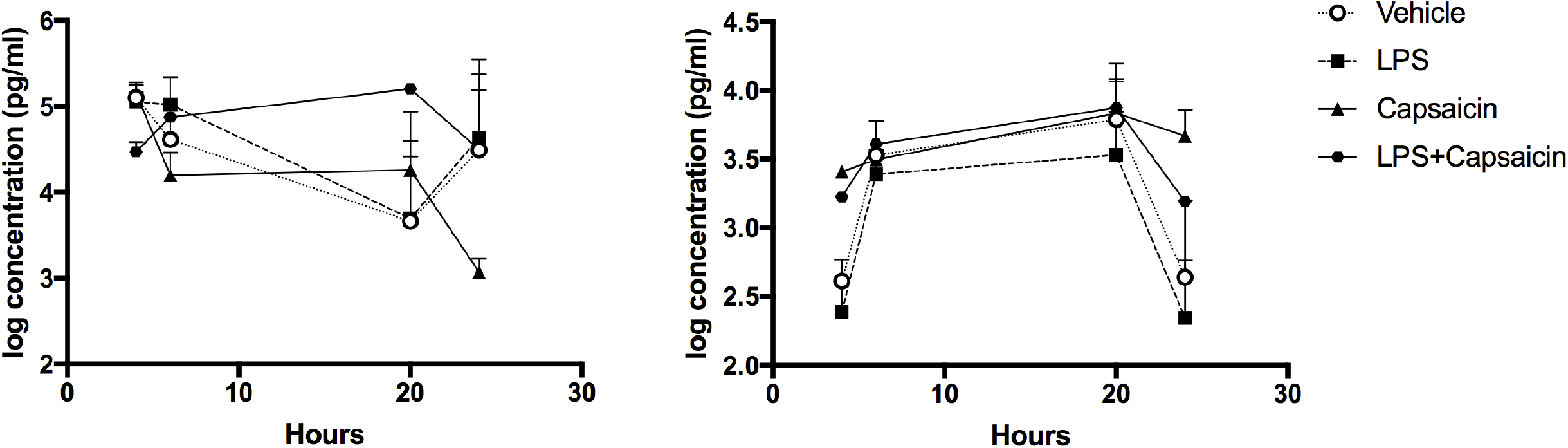

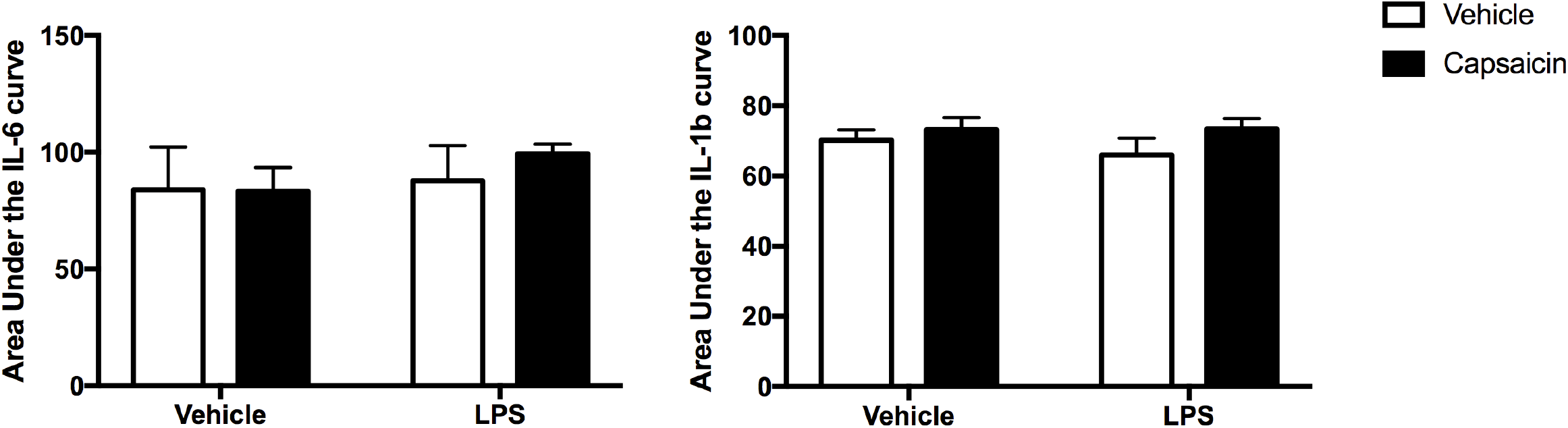

### LPS + capsaicin has an impact over time on stimulated IL-1β release from vaginal tissue sections

A three-way ANOVA of IL-1β release into the supernatant revealed a significant effect of stimulation time (*F*_3,32_=10.82, p<0.0001), LPS (*F*_1,32_=16.37, p=0.0003), but not capsaicin (*F*_1,32_=2, p>0.05), nor an interaction between the three (*F*_3,32_=0.1877, p>0.05)(Figure 1b). Post-hoc comparisons using Dunnett’s multiple comparisons test revealed significant differences between the 4-hour vehicle group and: 1) the 20-hour vehicle group (p<0.03); 2) the 20-hour capsaicin group (p<0.02); and 3) the 20-hour LPS+capsaicin group (p<0.02). The two-way ANOVA on IL-1β AUC data revealed a significant effect of capsaicin (*F*_1,8_=6.223, p=0.0373), but not LPS (*F*_1,8_=1, p>0.05), nor an interaction between the two (*F*_1,8_=1.095, p>0.05)(Figure 2b). Post-hoc comparisons using Sidak’s multiple comparisons test revealed no significant between-group differences.

## Discussion

This study has demonstrated that *ex vivo* stimulation of mouse vaginal tissues evokes a pro-inflammatory neuroimmune response. This response is evident as an increased IL-1β output following stimulation with the TRPV1 agonist, capsaicin; however, we observed no differences in IL-6 output. The optimal timeframe for *ex vivo* stimulation of mouse vaginal tissues to evoke an IL-1β response was 20 hours (Figure 1). To our knowledge, this is the first study to examine *ex vivo* stimulation of mouse vaginal tissues using both LPS and capsaicin.

Surprisingly, our study demonstrated higher mean-rank values of IL-1β output concentration following stimulation with capsaicin alone and a combination of LPS and capsaicin, rather than LPS alone. These findings suggest that capsaicin is driving a pro-inflammatory IL-1β response, and stimulation with LPS is contributing to the resolution of this pro-inflammatory response. This may be due to the regulatory effect that TLR4 activation has on cells that resolve inflammation. For example, the activation of TLR4 through LPS is associated with the release of neutrophil survival factors^42^, increasing the activation and survival time of neutrophils in a damaged area. Leukocytes such as neutrophils and eosinophils are essential in neutralising and eliminating pro-inflammatory mediators such as IL-1β^43^. This process of leukocytes eliminating pro-inflammatory mediators is necessary for the maintenance of homeostasis, and generally occurs before macrophages have migrated to the damaged area^43^. It is plausible to suggest that in the 24 hour period we observed, only inflammation-resolving cells such as leukocytes were activated. Due to the *ex vivo* nature of our work, there may have been insufficient macrophages in the tissue to produce a pro-inflammatory response in the later time frame.

Our study findings corroborate previous demonstrations that mouse female reproductive tract (FRT) tissues contain TLR4^44-46^ and TRPV1^47^. However, previous studies have demonstrated a pro-inflammatory response of IL-6 from uterine^45^ and endometrial^46^ cells following stimulation with LPS. Sheldon and Roberts (2010)^46^ observed the greatest output of IL-6 at 30 hours post-stimulation, which may explain why we did not observe an IL-6 response in our current study – perhaps we did not observe for long enough. No studies have investigated the response of IL-1β following stimulation of the mouse FRT with LPS. Only one study has investigated the expression of TRPV1 in the mouse FRT.^47^ De Clercq et al. (2017)^47^ used capsaicin to stimulate TRPV1 in uterine tissues, but expression levels of the responses following TRPV1 activation was below the level of detection post-stimulation. De Clercq et al. (2017)^47^ did not explore cytokine output following the stimulation with capsaicin. Immunohistochemistry findings reported by Barry et a. (2017)^48^ show a high number of substance P and calcitonin-gene related peptide (CGRP) positive nerve fibres in the mouse vagina, suggesting TRPV1 positive nerves are likely present. These findings, taken into consideration with our own, infer that TRPV1 is expressed in mouse vaginal tissues, but that is yet to be explored. Clearly more research into the functional expression of both TLR4 and TRPV1 and their response following stimulation in mouse vaginal tissues is warranted to further understand potential vaginal immune defense mechanisms.

The outcomes of this study relate specifically to mice but may also provide useful insight into a similar protocol for use in humans. Mouse models have provided tremendous insight into the workings of the innate and adaptive immune system in humans. Similar to the mouse FRT, human FRT tissues also express TLR4, although this expression decreases distally throughout the tract, and there are conflicting reports as to the presence or absence of TLR4 in the human vagina^49-51^. Stimulation of human uterine tissues with LPS has been shown to induce IL-8 output, but not IL-6 output^52^ or INF-γ^53^. No studies have investigated IL-1β output following stimulation of human FRT tissues with LPS. In humans, TRPV1 is present in the cervix^54^ and vulva vestibule^26^ of women. However, the rest of the human FRT has not been assessed for the presence of TRPV1, nor has the response of the FRT to stimulation with capsaicin been investigated. While there are differences between mouse and human immune system activation^55^, the similarities across species can provide useful insight into the immune function of the human vagina. Our model of stimulation with LPS and capsaicin should first be replicated in healthy human samples before it is used to investigate disease states such as vulvodynia.

Future research should consider limitations of our model. First, we have used tissue *ex vivo* for this model. The act of removing tissue will itself generate inflammation^56^, which may obscure a true measure of the LPS/capsaicin stimulations. Including a control vehicle stimulation aimed to account for this effect. Second, we only assessed the output of two cytokines, IL-1β and IL-6. The activation of TLR4 through stimulation with LPS has consistently shown to increase the output of TNF-α in mice in several studies using different tissues (for example, ^57-60^). We may have observed a different response if we assessed other cytokines such as TNF-α. Third, we did not assess the viability of cells at different time points nor perform a dose-response analysis. Future research should further explore the current model with greater sample sizes, inclusive of a broader range of cytokines, time points, and LPS and capsaicin doses. These limitations need to be addressed before firm conclusions can be drawn on the exact pattern of inflammatory response in this model.

In summary, we have demonstrated a pro-inflammatory neuroimmune response following *ex vivo* stimulation of mouse vaginal tissues with capsaicin and LPS, evident as an increased IL1-β output. The optimal timeframe for this stimulation is 20 hours. This model of *ex vivo* stimulation may be an effective method of investigating the neuroimmune interface in peripheral tissues. The 20 hours timeline may guide the use of this model in future work, particularly in the application to human models. Future research should aim to investigate the wider immune response generated by LPS and capsaicin in the mouse vagina and the mechanisms driving this response before the model is applied to human gynaecological disorders such as vulvodynia.

## References

1. Grace PM, Hutchinson MR, Maier SF, Watkins LR. Pathological pain and the neuroimmune interface. Nature Reviews Immunology. 2014;14(4):217–231.

2. Austin PJ, Moalem-Taylor G. The neuro-immune balance in neuropathic pain: involvement of inflammatory immune cells, immune-like glial cells and cytokines. Journal of neuroimmunology. 2010;229(1):26–50.

3. Grace PM, Rolan PE, Hutchinson MR. Peripheral immune contributions to the maintenance of central glial activation underlying neuropathic pain. Brain, behavior, and immunity. 2011;25(7):1322–1332.

4. Liu T, van Rooijen N, Tracey DJ. Depletion of macrophages reduces axonal degeneration and hyperalgesia following nerve injury. Pain. 2000;86(1):25–32.

5. Zuo Y, Perkins NM, Tracey DJ, Geczy CL. Inflammation and hyperalgesia induced by nerve injury in the rat: a key role of mast cells. Pain. 2003;105(3):467–479.

6. Perkins N, Tracey D. Hyperalgesia due to nerve injury: role of neutrophils. Neuroscience. 2000;101(3):745–757.

7. Kim D, Kim MA, Cho I-H, et al. A critical role of toll-like receptor 2 in nerve injury-induced spinal cord glial cell activation and pain hypersensitivity. Journal of Biological Chemistry. 2007;282(20):14975–14983.

8. Colburn R, DeLeo J, Rickman A, Yeager M, Kwon P, Hickey W. Dissociation of microglial activation and neuropathic pain behaviors following peripheral nerve injury in the rat. Journal of neuroimmunology. 1997;79(2):163–175.

9. Milligan ED, Watkins LR. Pathological and protective roles of glia in chronic pain. Nature reviews neuroscience. 2009;10(1):23–36.

10. Kawasaki Y, Zhang L, Cheng J-K, Ji R-R. Cytokine mechanisms of central sensitization: distinct and overlapping role of interleukin-1β, interleukin-6, and tumor necrosis factor-α in regulating synaptic and neuronal activity in the superficial spinal cord. Journal of Neuroscience. 2008;28(20):5189–5194.

11. Lehnardt S, Massillon L, Follett P, et al. Activation of innate immunity in the CNS triggers neurodegeneration through a Toll-like receptor 4-dependent pathway. Proceedings of the National Academy of Sciences. 2003;100(14):8514–8519.

12. Tang S-C, Arumugam TV, Xu X, et al. Pivotal role for neuronal Toll-like receptors in ischemic brain injury and functional deficits. Proceedings of the National Academy of Sciences. 2007;104(34):13798–13803.

13. Leon CG, Tory R, Jia J, Sivak O, Wasan KM. Discovery and development of toll-like receptor 4 (TLR4) antagonists: a new paradigm for treating sepsis and other diseases. Pharmaceutical research. 2008;25(8):1751–1761.

14. Nicotra L, Loram LC, Watkins LR, Hutchinson MR. Toll-like receptors in chronic pain. Experimental Neurology. 2012;234(2):316–329.

15. Diogenes A, Ferraz CCR, Akopian ANa, Henry MA, Hargreaves KM. LPS sensitizes TRPV1 via activation of TLR4 in trigeminal sensory neurons. Journal of dental research. 2011;90(6):759–764.

16. Hutchinson MR, Buijs M, Tuke J, et al. Low-dose endotoxin potentiates capsaicin-induced pain in man: evidence for a pain neuroimmune connection. Brain, behavior, and immunity. 2013;30:3–11.

17. Raetz CR, Whitfield C. Lipopolysaccharide endotoxins. Annual review of biochemistry. 2002;71(1):635–700.

18. Nagy J, Vincent S, Staines W, Fibinger H, Reisine T, Yamamura H. Neurotoxic action of capsaicin on spinal substance P neurons. Brain research. 1980;186(2):435–444.

19. Wharton J, Gulbenkian S, Mulderry P, et al. Capsaicin induces a depletion of calcitonin gene-related peptide (CGRP)-immunoreactive nerves in the cardiovascular system of the guinea pig and rat. Journal of the autonomic nervous system. 1986;16(4):289–309.

20. Caterina MJ, Schumacher MA, Tominaga M, Rosen TA, Levine JD, Julius D. The capsaicin receptor: a heat-activated ion channel in the pain pathway. Nature. 1997;389(6653):816–824.

21. Suffredini AF, Hochstein HD, McMahon FG. Dose-related inflammatory effects of intravenous endotoxin in humans: evaluation of a new clinical lot of Escherichia coli O: 113 endotoxin. Journal of Infectious Diseases. 1999;179(5):1278–1282.

22. Martich GD, Boujoukos AJ, Suffredini AF. Response of man to endotoxin. Immunobiology. 1993;187(3):403–416.

23. Chalmers KJ, Catley MJ, Evans SF, Moseley GL. Clinical assessment of the impact of pelvic pain on women. PAIN. 2016.

24. Harlow BL, Stewart EG. A population-based assessment of chronic unexplained vulvar pain: have we underestimated the prevalence of vulvodynia? Journal of the American Medical Women’s Association (1972). 2003;58(2):82–88.

25. Tympanidis P, Terenghi G, Dowd P. Increased innervation of the vulval vestibule in patients with vulvodynia. British Journal of Dermatology. 2003;148(5):1021–1027.

26. Tympanidis PC, M. A.; Yiangou, Y.; Terenghi, G.; Dowd, P.; Anand, P. Increased vanilloid receptor VR1 innervation in vulvodynia. European Journal of Pain. 2004;8 (2):129–133.

27. Farmer MA, Taylor AM, Bailey AL, et al. Repeated vulvovaginal fungal infections cause persistent pain in a mouse model of vulvodynia. Science Translational Medicine. 2011;3(101):101ra191–101ra191.

28. Sandig H, Bulfone-Paus S. TLR signaling in mast cells: common and unique features. Frontiers in immunology. 2012;3:185.

29. Netea MG, Van der Graaf CA, Vonk AG, Verschueren I, Van der Meer JW, Kullberg BJ. The role of toll-like receptor (TLR) 2 and TLR4 in the host defense against disseminated candidiasis. Journal of Infectious Diseases. 2002;185(10):1483–1489.

30. Grozdanovic Z, Mayer B, Baumgarten HG, Brüning G. Nitric oxide synthase-containing nerve fibers and neurons in the genital tract of the female mouse. Cell and tissue research. 1994;275(2):355–360.

31. Fox JG, Barthold S, Davisson M, Newcomer CE, Quimby FW, Smith A. The Mouse in biomedical research: diseases. Vol 2: Academic Press; 2006.

32. Jiang J, Kelly KA. Isolation of Lymphocytes from Mouse Genital Tract Mucosa. JoVE (Journal of Visualized Experiments). 2012(67):e4391–e4391.

33. Wormley Jr F, Scott M, Luo W, Baker M, Chaiban J, Fidel Jr P. Evidence for a unique expression of CD4 on murine vaginal CD4+ cells. Immunology. 2000;100(3):300–308.

34. Nandi D, Allison J. Characterization of neutrophils and T lymphocytes associated with the murine vaginal epithelium. Regional immunology. 1992;5(6):332–338.

35. Sonoda Y, Mukaida N, Wang J-b, et al. Physiologic regulation of postovulatory neutrophil migration into vagina in mice by a CXC chemokine (s). The Journal of Immunology. 1998;160(12):6159–6165.

36. Caligioni CS. Assessing reproductive status/stages in mice. Current protocols in neuroscience / editorial board, Jacqueline N Crawley [et al]. 2009;Appendix 4:Appendix 4I.

37. Cason AM, Samuelsen CL, Berkley KJ. Estrous changes in vaginal nociception in a rat model of endometriosis. Hormones and behavior. 2003;44(2):123–131.

38. Ariyadi B, Isobe N, Yoshimura Y. Toll-like receptor signaling for the induction of mucin expression by lipopolysaccharide in the hen vagina. Poultry science. 2014;93(3):673–679.

39. Miller MF, Loch-Caruso R. Comparison of LPS-stimulated release of cytokines in punch versus transwell tissue culture systems of human gestational membranes. Reproductive Biology and Endocrinology. 2010;8(1):121.

40. Ferraz CCR, Henry MA, Hargreaves KM, Diogenes A. Lipopolysaccharide from Porphyromonas gingivalis sensitizes capsaicin-sensitive nociceptors. Journal of endodontics. 2011;37(1):45–48.

41. Benson S, Kattoor J, Wegner A, et al. Acute experimental endotoxemia induces visceral hypersensitivity and altered pain evaluation in healthy humans. PAIN®. 2012;153(4):794–799.

42. Sabroe I, Prince LR, Jones EC, et al. Selective roles for Toll-like receptor (TLR) 2 and TLR4 in the regulation of neutrophil activation and life span. The Journal of Immunology. 2003;170(10):5268–5275.

43. Serhan CN, Brain SD, Buckley CD, et al. Resolution of inflammation: state of the art, definitions and terms. The FASEB journal. 2007;21(2):325–332.

44. Soboll G, Schaefer TM, Wira CR. Effect of Toll-Like Receptor (TLR) Agonists on TLR and Microbicide Expression in Uterine and Vaginal Tissues of the Mouse. American Journal of Reproductive Immunology. 2006;55(6):434–446.

45. Soboll G, Shen L, Wira CR. Expression of Toll-Like Receptors (TLR) and Responsiveness to TLR Agonists by Polarized Mouse Uterine Epithelial Cells in Culture 1. Biology of reproduction. 2006;75(1):131–139.

46. Sheldon IM, Roberts MH. Toll-like receptor 4 mediates the response of epithelial and stromal cells to lipopolysaccharide in the endometrium. PLoS One. 2010;5(9):e12906.

47. De Clercq K, Van den Eynde C, Hennes A, Van Bree R, Voets T, Vriens J. The functional expression of transient receptor potential channels in the mouse endometrium. Human Reproduction. 2017.

48. Barry CM, Ji E, Sharma H, et al. Morphological and neurochemical differences in peptidergic nerve fibers of the mouse vagina. Journal of Comparative Neurology. 2017;525(10):2394–2410.

49. Fazeli A, Bruce C, Anumba D. Characterization of Toll-like receptors in the female reproductive tract in humans. Human Reproduction. 2005;20(5):1372–1378.

50. Pioli PA, Amiel E, Schaefer TM, Connolly JE, Wira CR, Guyre PM. Differential expression of Toll-like receptors 2 and 4 in tissues of the human female reproductive tract. Infection and Immunity. 2004;72(10):5799–5806.

51. Pivarcsi A, Nagy I, Koreck A, et al. Microbial compounds induce the expression of pro-inflammatory cytokines, chemokines and human β-defensin-2 in vaginal epithelial cells. Microbes and Infection. 2005;7(9):1117–1127.

52. Schaefer TM, Desouza K, Fahey JV, Beagley KW, Wira CR. Toll-like receptor (TLR) expression and TLR-mediated cytokine/chemokine production by human uterine epithelial cells. Immunology. 2004;112(3):428–436.

53. Eriksson M, Meadows SK, Basu S, Mselle TF, Wira CR, Sentman CL. TLRs mediate IFN-γ production by human uterine NK cells in endometrium. The Journal of Immunology. 2006;176(10):6219–6224.

54. Tingåker BK, Ekman-Ordeberg G, Facer P, Irestedt L, Anand P. Influence of pregnancy and labor on the occurrence of nerve fibers expressing the capsaicin receptor TRPV1 in human corpus and cervix uteri. Reproductive Biology and Endocrinology. 2008;6(1):8.

55. Mestas J, Hughes CC. Of mice and not men: differences between mouse and human immunology. The Journal of Immunology. 2004;172(5):2731–2738.

56. Broughton 2nd G, Janis JE, Attinger CE. The basic science of wound healing. Plastic and reconstructive surgery. 2006;117(7 Suppl):12S–34S.

57. Nagai Y, Akashi S, Nagafuku M, et al. Essential role of MD-2 in LPS responsiveness and TLR4 distribution. Nature immunology. 2002;3(7):667–672.

58. Hoshino K, Takeuchi O, Kawai T, et al. Cutting edge: Toll-like receptor 4 (TLR4)-deficient mice are hyporesponsive to lipopolysaccharide: evidence for TLR4 as the Lps gene product. The Journal of Immunology. 1999;162(7):3749–3752.

59. Gendron R, Nestel F, Lapp W, Baines M. Lipopolysaccharide-induced fetal resorption in mice is associated with the intrauterine production of tumour necrosis factor-alpha. Journal of reproduction and fertility. 1990;90(2):395–402.

60. Zhang X, Morrison DC. Lipopolysaccharide-induced selective priming effects on tumor necrosis factor alpha and nitric oxide production in mouse peritoneal macrophages. Journal of Experimental Medicine. 1993;177(2):511–516.

